# One-pot enzymatic synthesis of the sugar nucleotide CDP-ribitol

**DOI:** 10.64898/2026.05.22.727167

**Authors:** Saeed Akkad, Eva W. Wan, Alex Maddison, Jessica Rule Mcloughlin, Cindy Au-Yeung, Lloyd D. Murphy, Lianne I. Willems

## Abstract

CDP-ribitol serves as an essential donor for D-ribitol-5-phosphate incorporation into bacterial and mammalian glycoconjugates. Here, we demonstrate the one-pot enzymatic synthesis of CDP-ribitol from the readily available precursor ribitol. We also explore the synthesis of bioorthogonally tagged derivatives and other CDP conjugates, thereby providing valuable new tools for glycoscience research.

---

D-ribitol-5-phosphate (RboP) is a pentitol phosphate that is an integral component of glycopolymers in both bacterial and mammalian systems. In Gram-positive bacteria, RboP is assembled into polymeric chains named wall teichoic acids (WTAs), anionic cell surface glycopolymers that are anchored to the peptidoglycan layer and contribute approximately half the cell wall mass. Together with lipoteichoic acids (LTAs), WTAs play important roles in cell division and cell shape. More importantly, they play a crucial role in the virulence, cell adhesion, biofilm formation and antibiotic susceptibility of some medically relevant bacteria, including *Staphylococcus aureus* and *Streptococcus pneumoniae*.^1–3^ In some Gram-negative bacteria, such as the pathogen *Haemophilus influenzae* type b, the capsular polysaccharide that is responsible for virulence is comprised of repeating units of β-D-ribose-D-ribitol-5-phosphate.^4^ Interestingly, RboP was not known to exist in mammals until only a decade ago, when it was found to be present in a single O-linked glycan on the cell surface glycoprotein α-dystroglycan (α-DG), which acts as a receptor for laminins and other extracellular matrix proteins and is essential in maintaining membrane integrity.^5–8^ Defects in the biosynthesis of this glycan, including the incorporation of RboP, have been associated with an array of muscular dystrophies known collectively as α-dystroglycanopathies.^9,10^

In each of these systems, RboP is transferred onto the acceptor glycan by ribitol-5-phosphate transferases which use the nucleotide-activated form of ribitol, cytidine diphosphate (CDP)-ribitol (Scheme 1), as a universal donor.^7,8,11,12^ The availability of CDP-ribitol as a research tool is essential for further studies into the assembly of RboP-containing glycoconjugates and how these can be manipulated for (bio)medical benefit. For example, because of the essential roles WTAs play in virulence and in interacting with the immune system in *S. aureus*-mediated disease, there is significant interest in these glycopolymers as a target for novel antimicrobials and as vaccine antigens.^13,14^ In addition, supplementation with CDP-ribitol is being investigated as a therapeutic strategy for α-dystroglycanopathies caused by deficiencies in the formation of RboP-containing glycans.^15^ CDP-ribitol is however not commercially available. Therefore, facile methods for the synthesis and isolation of CDP-ribitol on a preparative scale are highly desirable.

Chemical synthesis of CDP-ribitol has been reported in two steps from D-ribose-5-phosphate^16^ or in seven steps from D-ribose,^8^ both involving a key P(V)-P(V) coupling step to generate CDP-ribose or CDP-ribitol. Because some of these reactions and the purification of phosphorylated intermediates can be challenging, enzymatic synthesis offers an attractive alternative. Several enzymatic and chemoenzymatic procedures have been reported,^5,17–20^ which use cytidylyltransferases from bacterial or human origin (e.g. *S. aureus* TarI,^18^ *H. influenzae* Bcs1,^21^ or human ISPD)^5,7^ to produce CDP-ribitol from RboP and CTP. The RboP substrate for these enzymatic reactions is not commercially available and thus needs to be synthesised. This is usually done by reduction of D-ribose-5-phosphate and involves a non-trivial multistep purification procedure.^5^

As part of our ongoing research into the labelling of RboP-containing glycans,^22^ we were interested in developing an efficient enzymatic method for CDP-ribitol synthesis that would avoid the need to isolate RboP. To achieve this, we designed a strategy from the readily available and cheaper non-phosphorylated pentitol sugar, ribitol (currently costing around £150 for 100 g compared to £100 for 1 g of D-ribose-5-phosphate) (Scheme 1).

**Scheme 1.**
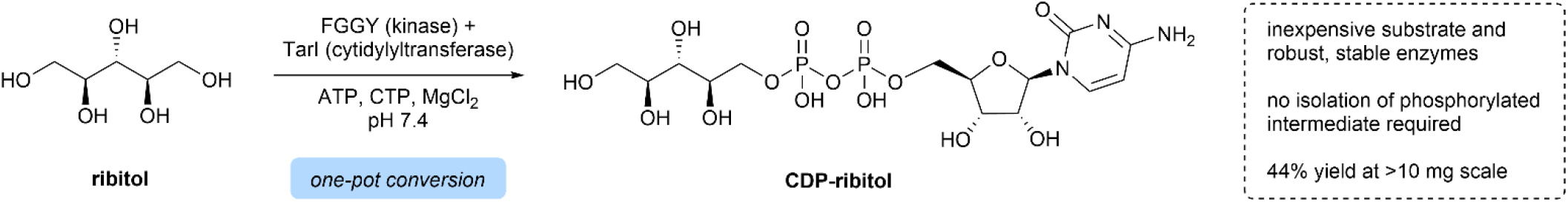
Enzymatic synthesis of CDP-ribitol via the one-pot conversion of the pentitol sugar ribitol using the human kinase FGGY and the cytidylyltransferase TarI from *S. aureus*.

Ribitol is not a precursor for CDP-ribitol biosynthesis in bacteria, which produce the donor by reduction of D-ribulose-5-phosphate, originating from the pentose phosphate pathway,^23^ to RboP and subsequent cytidylyl transfer to generate CDP-ribitol.^18,21^ In mammals, CDP-ribitol is similarly produced from RboP, but the metabolic origin of this substrate differs to that in bacteria. Recent evidence has revealed that several pathways are possible, in which RboP is formed either by reduction of D-ribose-5-phosphate, or via the reduction of D-ribose to ribitol.^24^ Notably, the latter suggests there must also be an enzyme that is able to phosphorylate the ribitol intermediate. Indeed, the human D-ribulose kinase FGGY was shown to display some activity on ribitol *in vitro*.^25^ Intrigued by the possibility of using this kinase as a strategy for RboP production, we set out to develop a one-pot enzymatic approach using FGGY together with the cytidylyltransferase TarI for the synthesis of CDP-ribitol (Scheme 1). Here, we describe the successful implementation of this novel methodology and demonstrate the isolation of the product CDP-ribitol on a multi-milligram scale. We also assess the promiscuity of the enzymatic system to synthesise unnatural, tagged derivatives of CDP-ribitol as potential bioorthogonal probes and expand its scope towards other biologically relevant CDP conjugates.

## Results and discussion

His-tagged human FGGY and His-tagged TarI from *Staphylococcus aureus* were readily produced by heterologous expression in an *E. coli* BL21 (DE3) expression system (see Supplementary Information (SI) for experimental procedures). The recombinant enzymes were purified by immobilized metal affinity chromatography and their activity was confirmed by monitoring product formation by LC-MS. Incubation of FGGY with ribitol and ATP led to the formation of a product at m/z 230.80 ([M-H]^-^), consistent with the mass of RboP and thus confirming the ability of the enzyme to phosphorylate ribitol (Figure S1).^25^ Additionally, using a pure sample of chemically synthesised RboP^26^ and CTP, TarI was shown to produce the expected CDP-ribitol product (m/z 536.06 ([M-H]^-^)) (Figure S2). We did not observe any loss in activity of either enzyme after storage of the purified aliquots for 12 months at -40 °C, suggesting the proteins are stable during long term storage.

Next, the two enzymatic reactions were combined to synthesise CDP-ribitol directly from ribitol. Ribitol was incubated with both enzymes along with ATP and CTP overnight, after which the mixture was treated with alkaline phosphatase (ALP) to dephosphorylate the free nucleotides,^5,7^ which interfered with the detection of CDP-ribitol in our LC-MS method due to peak overlap. Incidentally, ALP treatment also removes any remaining RboP intermediate, simplifying downstream purification.^17^ Upon quenching the reaction by heat inactivation, LC-MS analysis revealed a peak corresponding to CDP-ribitol, demonstrating that both enzymes acted consecutively to convert ribitol into the CDP-activated donor (Figure 1). The product was absent in control samples where one or both of the enzymes were omitted.

**Figure 1.**
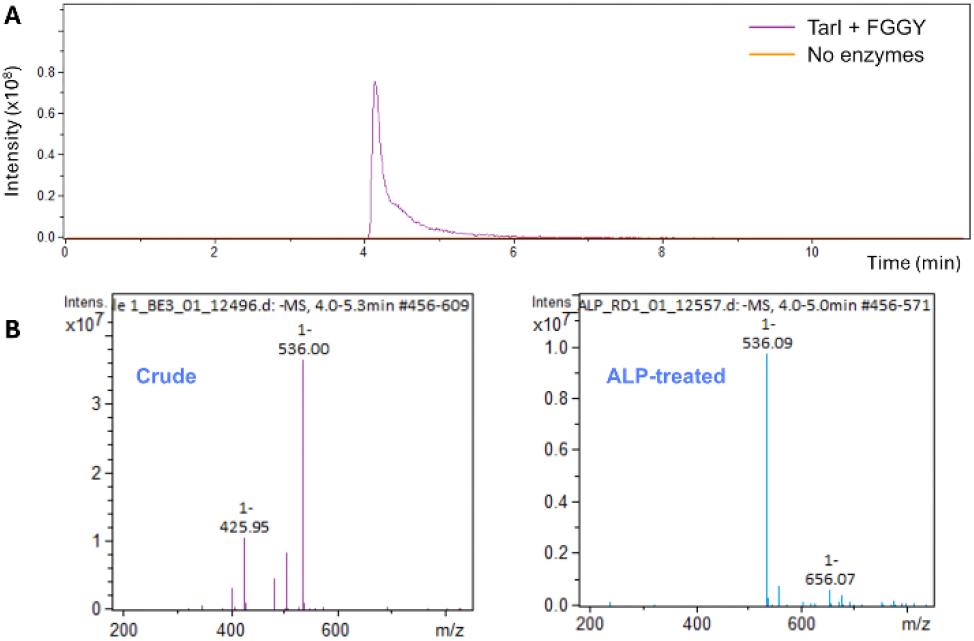
LC-MS analysis of CDP-ribitol formation after overnight incubation of ribitol with recombinant human FGGY and *S. aureus* TarI. A) Extracted ion chromatogram of CDP-ribitol (purple) compared to a non-enzyme treated control (orange). B) Mass spectrum of the enzyme-treated sample in (A) before and after ALP treatment, showing the formation of CDP-ribitol (m/z 536.09 ([M-H]^-^)).

Having shown that the two reactions successfully proceeded in one pot, we explored the utility of the new method to produce CDP-ribitol on a preparative scale. Hence, multiple 1 mL reactions, each containing 10 µmol of ribitol, were performed in parallel, ALP-treated, quenched and then pooled. The product was purified by anion exchange purification followed by reverse phase preparative HPLC to give pure CDP-ribitol, as confirmed by MS (Figure S3A) and NMR analysis (see SI for NMR spectra) in a yield of 17 mg (44%). Because issues with CDP-ribitol instability are described in the literature,^12,20,16^ we assessed the stability of the product in buffers of pH 4.5, 7.5 and 9.0 at 20, 37 or 60 °C overnight (Figure S3B). Despite small fluctuations in the concentration of the nucleotide sugar between samples, there was no consistent trend that would suggest significant loss of the product in the conditions tested. Additionally, we did not observe any notable degradation of the purified material during long-term storage at -20 °C. Thus, our approach enables the efficient production of CDP-ribitol on a multi-milligram scale without any significant stability issues.

We have recently reported the metabolic labelling of O-glycans on the human cell-surface protein α-dystroglycan with the use of alkyne-tagged RboP derivatives as glycan labelling probes.^22^ Because these probes require metabolic conversion into their corresponding CDP-activated donors before they can be incorporated into glycans, we were interested in exploring the synthesis of the bioorthogonally tagged CDP-ribitol derivatives using our new dual enzyme system. Therefore, two alkyne-tagged ribitol derivatives (probes A and B, Figure 2) were used as substrates in the one-pot dual enzyme reaction.

**Figure 2.**
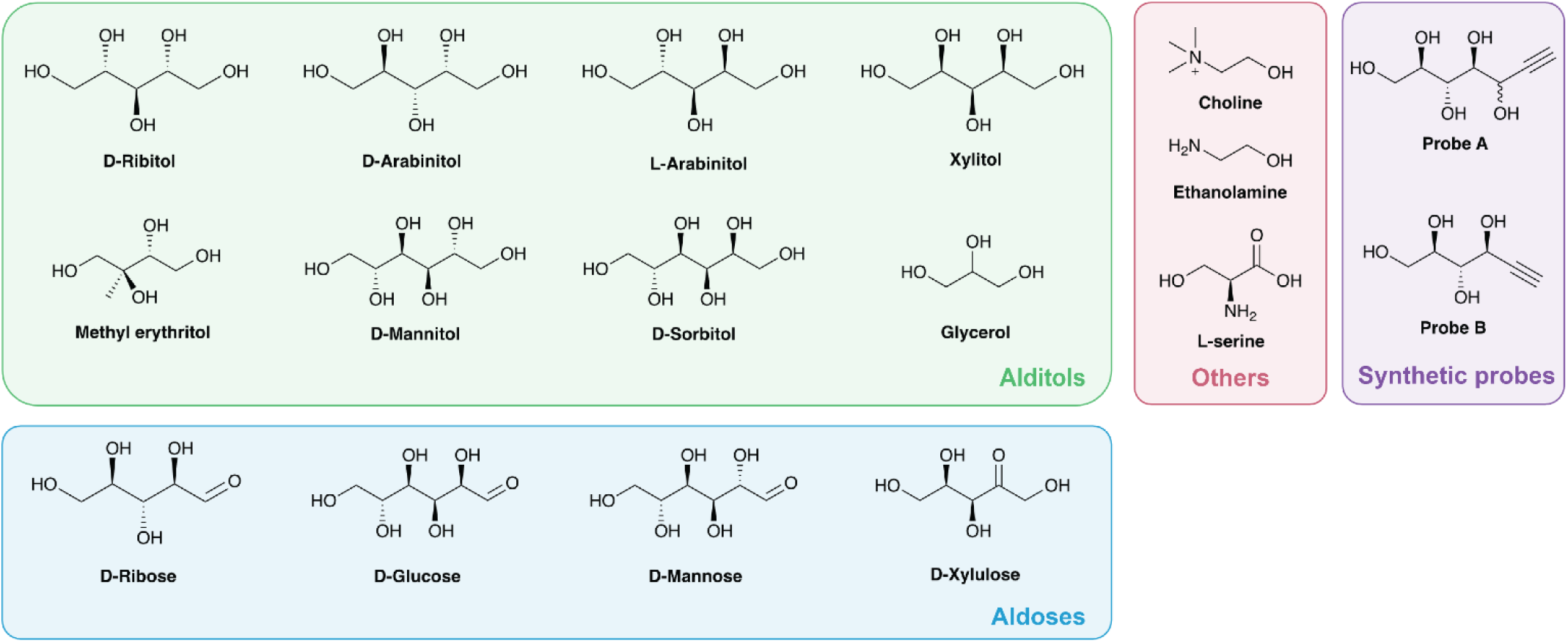
Structures of the substrates tested in the one-pot enzymatic reactions with FGGY/TarI and FGGY/PCYT2.

Conversion of probe A into the CDP-activated derivative was successful, although a higher substrate concentration was required for detectable product formation compared to non-tagged ribitol (Table 1, Figure S4). The CDP conjugate of the other tagged ribitol derivative, probe B, was not detected, but a peak corresponding to the phosphorylated intermediate was observed in the absence of ALP treatment (Figure S5). This suggests that the kinase displays higher substrate flexibility in the region around the C1-OH (numbering according to that in D-ribose) than the cytidylyltransferase. The ability to produce bioorthogonally tagged CDP-ribitol derivatives provides an interesting future avenue for the labelling of RboP-containing glycans, avoiding the need for cellular donor production and thereby potentially improving labelling efficiency compared to the use of alkyne-tagged RboP-derived probes.^22^ Although the nucleotide may cause issues with cell permeability, previous studies have shown that CDP-ribitol can be taken up by cells and enter the glycosylation pathway, supporting the feasibility of such a labelling strategy.

**Table 1.**
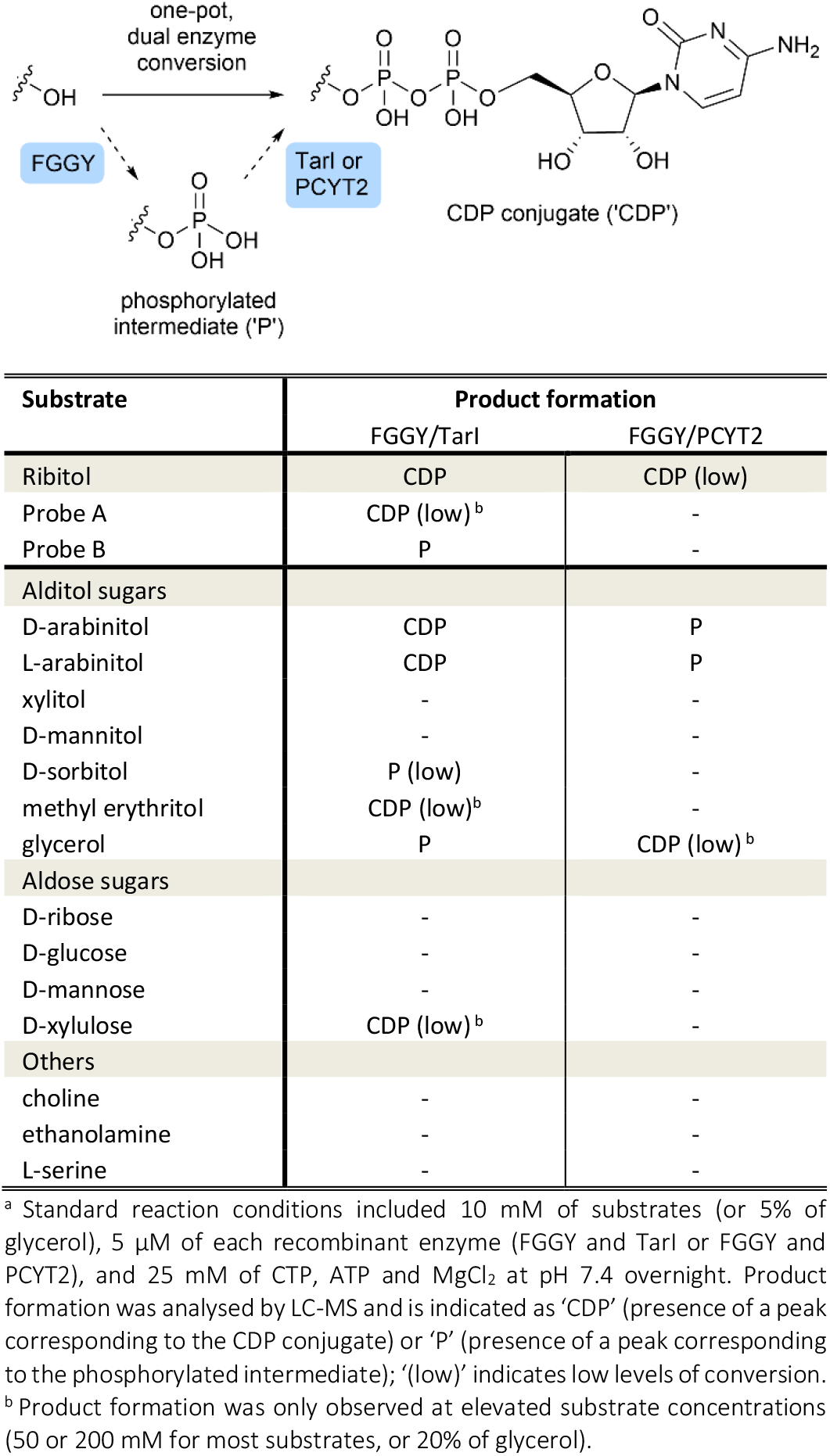
Synthesis of CDP-conjugates of various substrates by the enzyme pairs FGGY/TarI and FGGY/PCYT2^a^.

| Substrate | Product formation |  |
| --- | --- | --- |
|  | FGGY/TarI | FGGY/PCYT2 |
| Ribitol | CDP | CDP (low) |
| Probe A | CDP (low)<sup>b</sup> | - |
| Probe B | P | - |
| Alditol sugars |  |  |
| D-arabinitol | CDP | P |
| L-arabinitol | CDP | P |
| xylitol | - | - |
| D-mannitol | - | - |
| D-sorbitol | P (low) | - |
| methyl erythritol | CDP (low)<sup>b</sup> | - |
| glycerol | P | CDP (low)<sup>b</sup> |
| Aldose sugars |  |  |
| D-ribose | - | - |
| D-glucose | - | - |
| D-mannose | - | - |
| D-xylulose | CDP (low)<sup>b</sup> | - |
| Others |  |  |
| choline | - | - |
| ethanolamine | - | - |
| L-serine | - | - |

<sup>a</sup> Standard reaction conditions included 10 mM of substrates (or 5% of glycerol), 5 $\mu$ M of each recombinant enzyme (FGGY and TarI or FGGY and PCYT2), and 25 mM of CTP, ATP and MgCl<sub>2</sub> at pH 7.4 overnight. Product formation was analysed by LC-MS and is indicated as 'CDP' (presence of a peak corresponding to the CDP conjugate) or 'P' (presence of a peak corresponding to the phosphorylated intermediate); '(low)' indicates low levels of conversion.
<sup>b</sup> Product formation was only observed at elevated substrate concentrations (50 or 200 mM for most substrates, or 20% of glycerol).

Encouraged by the efficiency and the scalability of the approach, we next explored its substrate flexibility for the synthesis of other biologically relevant CDP conjugates and structurally similar analogues. Small scale reactions were performed using various substrates (Figure 2) and their conversion was monitored by LC-MS analysis (Table 1). Of the alditol sugars tested, both D-arabinitol and L-arabinitol showed conversion into their respective CDP conjugates (Table 1, Figure S6). D-arabinitol-1-phosphate is incorporated into the capsular polysaccharide of some serotypes of *S. pneumoniae* to evade the host immune response, with CDP-D-arabinitol acting as the donor.^27,28^ Interestingly, xylitol did not show any conversion, despite its structural similarity to both ribitol and arabinitol. This suggests the precise spatial relationship between the hydroxyl substituents is important in ensuring a productive fit in the active site of the enzymes. Surprisingly, the pentose analogue D-xylulose was converted into CDP-D-xylulose (Figure S7), which is a biosynthetic precursor to CDP-D-arabinitol in *S. pneumoniae*,^28^ though only at high substrate concentration.

Upon testing further alditol and aldose substrates, we did not detect the formation of CDP conjugates of D-sorbitol, D-mannitol (which is used in capsular polysaccharide biosynthesis),^29^ D-ribose, D-glucose (an intermediate in the biosynthesis of rare dideoxyhexoses for lipopolysaccharide biosynthesis)^30^ or D-mannose. Since FGGY was previously shown to phosphorylate D-ribose at similar levels to D-xylulose,^25^ the lack of nucleotide product formation from ribose suggests that TarI is unable to use ribose-phosphate as a substrate.

In contrast, the branched polyol substrate methyl erythritol was converted into CDP-methylerythritol when used at elevated substrate concentrations (Table 1, Figure S8). CDP-methylerythritol is an important intermediate in the 2C-methylerythritol 4-phosphate (MEP) pathway which is linked to the biosynthesis of isoprenoids in plants, algae, and bacteria. There is significant interest in its synthesis for the purpose of enzyme discovery, as a precursor for isoprenoid-derived natural products and as a target for the development of antimicrobials.^31,32^ While our method was able to produce CDP-activated methylerythritol, it should be noted that the small scale of these experiments prevented confirmation of the position at which the CDP group is attached to the polyol. Unlike the product of ribitol turnover by FGGY and TarI, which has been characterised previously, the exact structure of the products formed from other substrates remains to be determined.

The FGGY/TarI one-pot reaction was unable to convert the smaller substrates glycerol, ethanolamine and choline into their CDP-activated derivatives (Table 1). CDP-ethanolamine and CDP-choline are intermediates in the biosynthesis of membrane phospholipids.^33^ CDP-glycerol is used for the biosynthesis of both Gram-positive and Gram-negative bacterial glycopolymers, enabling the incorporation of *sn*-glycerol-3-phosphate into WTAs and capsular polysaccharides.^2,34^While the former two CDP conjugates are commercially available, CDP-glycerol is not. Access to this donor is important to facilitate enzyme discovery and the development of capsular polysaccharide-based vaccines.^34,35^ Enzymatic synthesis of CDP-glycerol has been described in the literature, starting from glycerol-phosphate^35,36^ or glycerol^37^ and using the glycerol-phosphate cytidylyltransferases Cps3B/Cps7B or TagD, along with the glycerol kinase GlpK. We found that the FGGY/TarI enzyme reaction produced a phosphorylated intermediate, indicating that FGGY was able to act on glycerol as a substrate. However, the CDP-activated donor was not detected, which is perhaps unsurprising given that *S. aureus* expresses distinct cytidylyltransferases to produce CDP-glycerol and CDP-ribitol (TarD and TarI, respectively). Given the importance of CDP-glycerol, we decided to expand our toolkit with the recently identified human cytidylyltransferase PCYT2, which is involved in the biosynthesis of the phospholipid phosphatidyl-ethanolamine but also showed activity on glycerol-phosphate.^38^ His-tagged PCYT2 was expressed and purified as above and then used in combination with FGGY for the one-pot conversion of glycerol. This was successful, generating the expected CDP-glycerol product at m/z 476.15 ([M-H]^-^) (Figure S9). Interestingly, the same enzyme combination was also able to generate CDP-ribitol, albeit at lower levels than the reaction containing FGGY and TarI.

## Conclusions

This work describes a novel strategy for the enzymatic synthesis of several biologically important CDP conjugates. In particular, we have demonstrated the efficient synthesis of CDP-ribitol via a dual-enzyme, one-pot conversion of the pentitol sugar, ribitol. Key to the success of our approach was the use of the human ribulokinase FGGY, which displays remarkable substrate flexibility. Coupled with the bacterial cytidylyltransferase TarI, our approach enables access to pure CDP-ribitol in multi-milligram quantities without specialist chemical synthesis expertise. This will facilitate studies aimed at improving our understanding of the biosynthetic pathways towards RboP-containing glycoconjugates and the characterisation of novel enzyme activities. Additionally, our work offers the potential to accelerate the development of emerging therapeutic approaches, such as CDP-ribitol supplementation for the treatment of α-dystroglycanopathies, WTA biosynthesis inhibitors as antimicrobial agents and synthetic glycoconjugate vaccines.^14,15^ The flexibility of the FGGY/TarI pair in generating bioorthogonally tagged CDP-ribitol derivatives also creates new opportunities in glycan engineering, expanding the available toolkit for glycoscience research.

## Experimental procedures

All experimental procedures, supporting figures, and NMR spectra of CDP-ribitol and new compounds can be found in the Supplementary Information file.

## Author contributions

S.A. investigation; methodology; validation; visualisation; data curation; writing - original draft. E.W.W., A.M., J.R.M. and C.A-Y. investigation. L.D.M. resources. L.I.W. conceptualisation; funding acquisition; writing - review & editing; visualisation; supervision; project administration.

## Conflicts of interest

There are no conflicts to declare.

## Acknowledgements

This work was supported by the project ‘RibiTool’ funded by the European Research Council (ERC) under the European Union’s Horizon 2020 research and innovation programme [Grant Agreement No 851448].

## Notes

### Competing Interest Statement

The authors have declared no competing interest.

