## Supplementary Information for "One-pot enzymatic synthesis of the sugar nucleotide CDP-ribitol"

**Experimental procedures** **2**
General methods 2
Chemical synthesis of ribitol-5-phosphate (RboP) barium salt 2
Synthesis of 1-ethynyl-D-ribitol (Probe A) 3
Synthesis of (2S,3S,4R)-2,3,4,5-tetrahydroxy-pent-1,6-yne (Probe B) 3
Expression and purification of recombinant enzymes 4
Enzymatic synthesis of CDP-ribitol 4
General method for the enzymatic synthesis of CDP-derivatives 5
LC-MS analysis of the enzymatic reaction products 6
LC-MS/MS quantitative analysis of CDP-Rbo 6
**Supporting figures** **7**
**NMR spectra** **12**

### Experimental procedures

**General methods**

All reactions were conducted in oven-dried glassware and dry $CH_2Cl_2$ was freshly distilled. $^1H$ and $^{13}C$ NMR spectra were recorded on a JEOL ECS400A spectrometer (400 and 101 MHz respectively). Structural assignments were corroborated by homo and heteronuclear 2D NMR methods (COSY, HMQC and DEPT) where necessary. Chemical shifts are reported in parts per million (ppm, δ) relative to the solvent. $^1H$ NMR splitting patterns are designated as singlet (s), doublet (d), triplet (t), quartet (q), doublet of doublets (dd), doublet of doublets of doublets (ddd), doublet of triplets (dt), or multiplet (m). Coupling constants (*J*) are reported in Hertz (Hz). The integrals reported in the $^1H$ NMR peak list for mixtures of diastereomers were normalised so that the numbers of protons for each diastereomer are reported in whole integers. The actual diastereomer ratio is also given and was determined from the relative integration of the respective proton peaks..

**Chemical synthesis of ribitol-5-phosphate (RboP) barium salt**

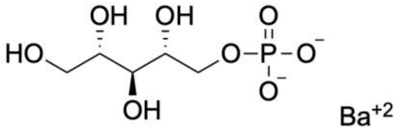


Method adapted from Baddiley et al.[39] 200 mg (0.42 mmol) of D-ribose-phosphate barium salt was dissolved in 5 mL of deionised water. The solution was cooled down in an ice bath and about 50 mg (1.32 mmol) of sodium borohydride was added slowly in batches. The reaction was allowed to reach room temperature before being stirred overnight. The mixture was cooled down again and quenched by the slow addition of 1 mL of glacial acetic acid. The pH of the solution was then adjusted to ~9 using dilute ammonia (1M).

5 g of AG1 X-8 resin (acetate form) was equilibrated with a solution of ammonium acetate (pH 8) with slight agitation for 30 min. The supernatant was then removed and the sample applied and left incubating with the resin (with agitation) for 1h. After removing the supernatant, the resin was washed twice with the ammonium acetate solution. 20 mL of ammonium formate 1N was then added to the resin and the mixture was left shaking for 1h to elute the product.

3 g of Amberlite IR-120 (H form) was washed three times with 15 mL of 4% HCl and then three times with deionised water (until the pH of the supernatant was neutral). The eluate from the AG1 X-8 purification was then added to the Amberlite resin and left shaking at room temperature for 1h, before being collected and filtered. The flow-through was lyophilised. To remove any traces of borate, the dry residue was redissolved in anhydrous methanol, stirred for 30 min, and the solvent was removed under reduced pressure. This process was repeated two more times. The dry residue was redissolved in deionised water, and 5 mL 0.1N barium hydroxide was added followed by a piece of dry ice. After the sample had set, any solid residue was removed by centrifugation. The supernatant was collected and an excess of ethanol was then used to precipitate RboP as its barium salt. The precipitate was washed with acetone and dried. **ESI-MS:** $[M-H]^-$ calculated for $C_5H_{12}O_8P$ 231.03, found 230.82. Characterisation was in accordance with the data reported in literature.[26,39]

**Synthesis of 1-ethynyl-D-ribitol (Probe A)**

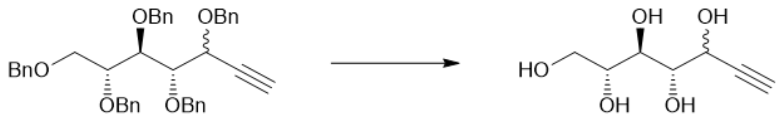


Fully benzylated 1-ethynyl-D-ribitol, described previously[22] (72.0 mg, 0.409 mmol, 1.0 equiv) was dissolved in anhydrous $CH_2Cl_2$ (3.8 mL) under $N_2$ atmosphere and cooled to –78 ℃. 1.0M $BCl_3$ in $CH_2Cl_2$ (3.8 mL, 3.80 mmol, 9.5 equiv) was added very slowly, and the mixture was stirred for 3 hours at –78℃. Upon completion the reaction was quenched with 20:1 MeOH/$H_2O$ and stirred for 30 minutes before warming to room temperature. The mixture was evaporated to dryness in vacuo. The sample was purified by reverse phase column chromatography using a C18 column on an ACCQPrep (10-80% $H_2O$/acetonitrile/0.1% formic acid) to obtain the product, Probe A, as a dark orange residue (22.8 mg, 0.130 mmol, 32%) in a 1.1:1 ratio of diastereoisomers. $R_f$ 0.56 (1:4 MeOH/$CH_2Cl_2$). HRMS (ESI): Calcd for $C_7H_{12}NaO_5$ $[M+Na]^+$ 199.0582, found 199.0577. $^1H$ NMR of isomeric mixture (400 MHz, DMSO-$D_6$) δ(ppm) 4.90 (d, J = 6.4 Hz, 1H, OH), 4.61 (d, J = 4.6 Hz, 1H, OH), 4.48 (t, J = 6.8 Hz, 2H, OH), 4.43 (dd, J = 5.0, 2.1 Hz, 1H), 4.35 (dq, J = 4.6, 2.3 Hz, 2H), 4.27 (dd, J = 8.0, 1.6 Hz, 1H, OH), 3.66 – 3.54 (m, 8H), 3.22 (dd, J = 8.7, 6.4 Hz, 1H, H7). $^{13}C$ NMR (101 MHz, DMSO-$D_6$) δ(ppm) 107.6 (C6-major), 106.9 (C6-minor), 77.2 (C1-major), 77.0 (C1-minor), 75.2 (C4-major), 73.5 (C4-minor), 70.2 (C7), 69.8, 69.3, 65.5, 65.0, 63.6.

**Synthesis of (2S,3S,4R)-2,3,4,5-tetrahydroxy-pent-1,6-yne (Probe B)**

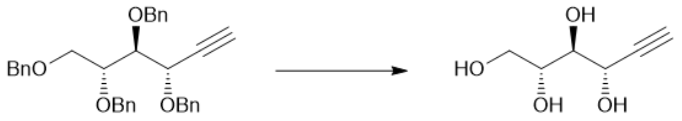


(2S,3S,4R)-2,3,4,5-tetrabenzyloxy-pent-1,6-yne, described previously[22] (180 mg, 350 μmol, 1.0 equiv) was dissolved in anhydrous $CH_2Cl_2$ (10 mL) under $N_2$ atmosphere and cooled to –78 ℃. 1.0M $BCl_3$ in $CH_2Cl_2$ (1.75 mL, 1.75 mmol, 5.0 equiv) was added slowly to the reaction. The mixture was stirred at –78 ℃ for 2 hours before warming to 0 ℃ over 3 hours. Upon completion the reaction was quenched with 20:1 MeOH/$H_2O$ and stirred for 30 minutes before warming to room temperature. HRMS (ESI) calculated for $C_6H_{10}KO_4$ $[M+K]^+$ 185.0216, found 185.0211. $^1H$ NMR (400 MHz, $CD_3OD$) δ(ppm) 4.56 – 4.55 (t, *J* = 3.1 Hz, 1H, H2), 3.76 (dd, *J* = 11.1, 2.7 Hz, 1H, H5), 3.68 – 3.56 (m, 3H, H3 + H4 + H5), 2.77 (d, *J* = 2.2 Hz, 1H, H6). $^{13}C$ NMR (101 MHz, $CD_3OD$) δ(ppm) 82.9 (C6), 75.4 (C4), 75.4 (C2), 73.6 (C3), 64.9 (C2), 64.5 (C5).

**Expression and purification of recombinant enzymes**

The sequence encoding TARI1_STAA8 from *S. aureus* was optimised for *E. coli* expression and ordered from Genscript.

Forward primer 5'-CCGCGCGGCAGCCATATGAAGTATGCGGGTATCCTGGC-3' and reverse primer 5'-AGTCATGCTAGCTTAGTCGTCCGCAATACCACCA-3' were used to clone the TarI gene into a pET-28a(+) vector using In-Fusion Cloning kit (TakaraBio). The ligation product was confirmed using Next-Gen Whole Plasmid Sequencing (Eurofins).

The sequences encoding human FGGY and PCYT2 were optimised for bacterial expression and ordered from Genscript in pET-28a(+).

Expression conditions used for all enzymes were identical: 5 ng of the plasmid was used to transform BL21(DE3)-Gold chemically competent cells (New England Biolabs) by heat shock and selected with kanamycin (50 µg/mL) at 37 °C for 16 h. Starter cultures were prepared by picking single colonies into LB with kanamycin (50 µg/mL) and grown at 37 °C for 16 h with shaking (180 rpm). Flasks containing LB with kanamycin (50 µg/mL) were inoculated with the starter culture at a dilution of 1:100 and grown at 37 °C with shaking (180 rpm) until OD600 reached ~0.6. At that point the cultures were induced with isopropyl β-D-1-thiogalactopyranoside (IPTG) to a final concentration of 1 mM, then the cultures were maintained shaking (180 rpm) at 18 °C overnight.

For the purification of the enzymes, the cells were harvested by centrifugation (6000 xg, 4 °C, for 10 min) and the pellets were resuspended in lysis buffer (25 mM Tris HCl, NaCl 300 mM, imidazole 5 mM, pH 7.8, protease inhibitor cocktail). The cells were lysed by sonication, centrifuged at 27,000 xg, 4 °C, for 40 min, and the enzymes were purified from their respective lysates by IMAC using a HisTrap FF Crude 5 mL column (Cytiva) and a gradient of elution buffer (25 mM Tris HCl, NaCl 300 mM, imidazole 500 mM). The eluted fractions were analysed by SDS-PAGE, and the fractions containing the target enzyme were pooled and desalted on a HiPrep G-25 desalting column (Cytiva) into storage buffer (25 mM Tris HCl, NaCl 150 mM, pH 7.8). The enzymes were stable in that buffer and could be aliquoted and stored in the -40 °C freezer, except for His-PCYT2, which required the addition of 20% glycerol to stabilise it. Since glycerol would interfere with the downstream applications of PCYT2, before each enzymatic reaction the thawed aliquots of the enzyme were desalted on a 5mL HiTrap G-25 desalting column (Cytiva) into a glycerol-free buffer and used immediately.

**Enzymatic synthesis of CDP-ribitol**

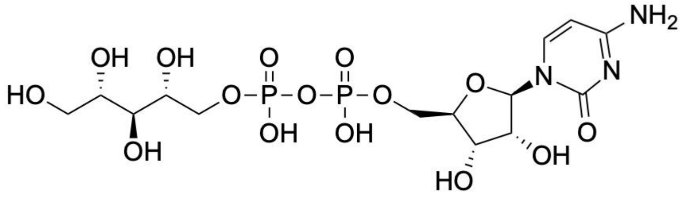


Note: ATP and CTP were prepared as stock solutions at a concentration of 500 mM buffered with Tris base to a pH between 7.3 and 7.5. We found this step was important since using unbuffered stock solutions resulted in acidification of the reaction mixture and caused precipitation of the proteins.

Seven identical reactions were set up in Eppendorf tubes, each containing ribitol at a final concentration of 10 mM (0.010 mmol per reaction), ATP, CTP, and $MgCl_2$ at a final concentration of 25 mM each, and His-FGGY and His-TarI at a final concentration of 5 µM each in a total volume of 1 mL of reaction buffer (Tris HCl 25 mM, NaCl 150 mM, pH 7.4). The tubes were incubated in a thermal shaker at 37 ℃/600 rpm overnight. The reaction was stopped by boiling the samples at 95 ℃ for 10 min. The samples were then spun down and the pellet discarded. To each sample was added 25 units of FastAP alkaline phosphatase, along the supplier-provided reaction buffer, and the samples were incubated in a thermal shaker at 37 ℃/600 rpm overnight. The reaction was stopped by boiling the samples at 95 ℃ for 10 min then the samples were spun down and the pellet discarded. Small aliquots were used to confirm the disappearance of the nucleotides from the reaction mixture by LC-MS, before the samples were pooled and run through a 3 kDa MWCO spin concentrator to remove any remaining protein. The flow-through was collected and its pH adjusted to ~9 with dilute ammonia (1M). The sample was then added to 3 g of AG-1 x8 resin (acetate form) and gently agitated at room temperature for 1h. The supernatant was discarded and the resin was washed twice with deionised water. The product was eluted with 20 mL of 1N ammonium formate, after which the eluate was lyophilised.

The dry residue was weighed to determine the loading for the prep HPLC purification. The sample was then dissolved in an appropriate volume of HPLC-grade water containing 0.1% formic acid and was purified on a Teledyne ACCQprep H150 prep HPLC system using a C18AQ RediSep column (30x150 m, 5 µm) as a stationary phase and water/acetonitrile (each containing 0.1% FA) as a mobile phase. The run was set at an isocratic step of 100% water (0.1% FA) at a flow rate 42.5 mL/min for 3 column volumes (CVs), followed by a stepped gradient of acetonitrile (0.1% FA) for 4 CVs, and finally an isocratic step of 100% acetonitrile (0.1% FA) for 2 CVs. Peak fractions were analysed by LC-MS, and the target compound was found to have eluted in the first isocratic step of 100% water. The fractions containing the product were pooled and lyophilised to give CDP-ribitol in a yield of 16.6 mg (0.031 mmol, 44%). ESI-MS: $[M-H]^-$: calculated for $C_{14}H_{24}N_3O_{15}P_2$ 536.08; found 536.04. $^1H$ NMR (400 MHz, $D_2O$): δ(ppm) 8.11 (d, $J$ = 7.9 Hz, 1H), 6.23 (d, $J$ = 7.8 Hz, 1H), 5.92 (d, $J$ = 3.2 Hz, 1H), 4.23-4.235 (m, 4H), 3.89-4.21 (m, 3H), 3.68–3.93 (m, 4H), 3.65-3.65 (m, 1H). $^{13}C$ NMR (101 MHz, $D_2O$) δ(ppm) 169.81, 161.22, 143.36, 95.69, 89.49, 83.11 (d, $J$ = 8.7 Hz), 74.35, 72.08, 71.62, 70.86 (d, $J$ = 7.8 Hz), 69.16, 67.01 (d, $J$ = 5.8 Hz), 64.47, 62.33. $^{31}P$ NMR (162 MHz, $D_2O$) δ 10.27 (d, $J$ = 112.0 Hz).

**General method for the enzymatic synthesis of CDP-derivatives**

ATP and CTP were prepared as stock solutions at a concentration of 500 mM buffered with Tris base to a pH between 7.3 and 7.5.

To a reaction mixture (final volume 100 µL) containing ATP, CTP and $MgCl_2$ each at a final concentration of 25 mM in reaction buffer (Tris HCl 25 mM, NaCl 150 mM, pH 7.4) and the kinase and cytidylyltransferase both at a final concentration of 5 µM, substrate was added at a final concentration of 10 mM, unless indicated otherwise. The reaction tube was incubated in a thermal shaker at 37 ℃/600 rpm overnight. The reaction was then stopped by boiling the sample at 95 ℃ for 10 min. The sample was then spun down and the pellet discarded. The supernatant was used to analyse product formation by LC-MS. Quantitative analysis could not be performed due to the lack of pure products to use as standards, but relative peak intensities were used to qualitatively assess the level of conversion.

Individual FGGY and TarI reactions to validate enzyme activity were performed essentially as above, but using ribitol as a substrate with FGGY and ATP (without TarI/CTP), or synthetic RboP as a substrate with TarI and CTP (without FGGY/ATP). Control reactions were performed without enzyme or with enzyme that had been denatured by boiling at 95 ℃ for 10 min.

**LC-MS analysis of the enzymatic reaction products**

High Performance Liquid Chromatography Electrospray Ionisation Mass Spectrometry (HPLC-MS) analysis of the CDP derivatives was performed using a Dionex UltiMate® 3000 Ci Rapid Separation LC system equipped with an UltiMate® 3000 photodiode array detector probing at 250–400 nm, coupled to a HCT ultra ETD II (Bruker Daltonics) ion trap spectrometer, using Chromeleon® 6.80 SR12 software (ThermoScientific), esquireControl version 6.2, Build 62.24 software (Bruker Daltonics), and Bruker compass HyStar 3.2-SR2, HyStar version 3.2, Build 44 software (Bruker Daltonics). All mass spectrometry was conducted in negative ion mode. Data analysis was performed using ESI Compass 1.3 DataAnalysis, Version 4.1 software (Bruker Daltonics). Samples were chromatographically analysed using an Accucore HILIC column, 2.6 µ (50 mm × 2.1 mm, Thermoscientific). Samples from the reaction mixtures were run crude without any further treatment. Water, 0.1% formic acid by volume (solvent A), and acetonitrile, 0.1 % formic acid (solvent B) were used as the mobile phase at a flow rate of 0.3 $mL.min^{-1}$ at 30 °C. A 1-minute step with 95% solvent B was used to load the sample onto the column, then a gradient from 95% to 5% solvent B over 5 minutes was used to elute the target compound. The mobile phase was held at 5% solvent B for 2.5 minutes before shifting it back to 95% and equilibrating the column again at that concentration for 2 minutes for the next run.

**LC-MS/MS quantitative analysis of CDP-Rbo**

High Performance Liquid Chromatography-Tandem Mass Spectrometry (HPLC-MS/MS) was performed on a Dionex UltiMate 3000 UHPLC system (ThermoScientific) coupled to a TSQ Endura triple quadrupole mass spectrometer (ThermoScientific) via a heated electrospray ionisation (H-ESI) source operated in negative ion mode. For the standard calibration curve, a series of standard solutions with known concentrations of pure CDP-Rbo were prepared in LC-MS grade water then analysed. For analysing samples from the stability study, an aliquot was taken from the incubated solution, diluted with LC-MS grade water at a ratio of 1:1, then analysed. Sample analysis was performed using an Accucore HILIC column, 2.6 µ (50 mm × 2.1 mm, Thermoscientific). Samples from the reaction mixtures were run crude without any further treatment. Water, 0.1% formic acid by volume (solvent A), and acetonitrile, 0.1 % formic acid (solvent B) were used as the mobile phase at a flow rate of 0.3 $mL.min^{-1}$ at 30 °C. A 1-minute step with 95% solvent B was used to load the sample onto the column, then a gradient from 95% to 5% solvent B over 5 minutes was used to elute the target compound. The mobile phase was held at 5% solvent B for 2.5 minutes before shifting it back to 95% and equilibrating the column again at that concentration for 2 minutes for the next run. Mass spectrometric detection was performed using the following source parameters: spray voltage, 2500 V; sheath gas, 50 (arbitrary units); auxiliary gas, 10; sweep gas, 1; ion transfer tube temperature, 325 °C; and vaporiser temperature, 350 °C. CDP-ribitol was quantified by selected reaction monitoring (SRM) of the transition m/z 536.038 to 321.982, with a collision energy of 20.33 V and an RF lens voltage of 184 V, acquired between 0.5 and 12.0 min. Q1 and Q3 resolutions were set to 0.7 and 1.2 Da (FWHM), respectively. Data were acquired and processed using Xcalibur version 4.0.27.10 (ThermoScientific).

#### Supporting figures

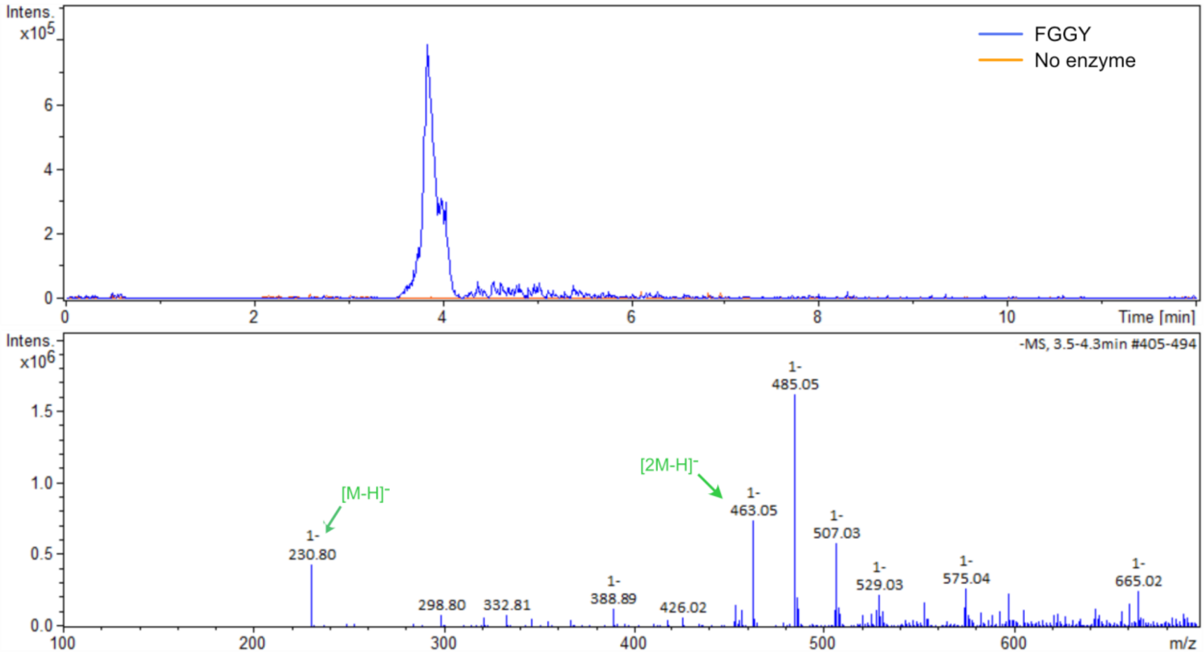


**Figure S1.** Top: Extracted ion chromatogram (EIC) of RboP from a ribitol sample treated with FGGY compared to an untreated (no enzyme) control. Bottom: mass spectrum from the sample treated with the active enzyme, showing the formation of RboP (m/z 230.80 ([M-H]-).

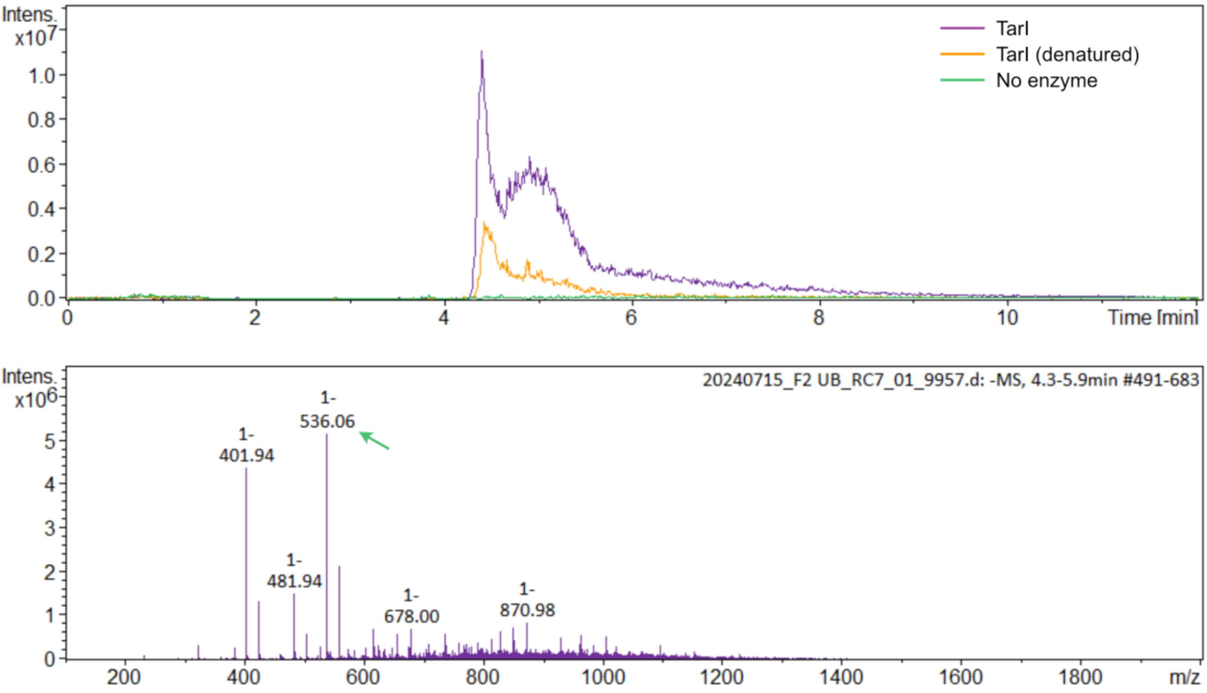


**Figure S2.** Top: Extracted ion chromatogram (EIC) of CDP-Rbo from a sample of chemically synthesised RboP treated with TarI compared to an untreated (no enzyme) control and a control treated with denatured enzyme. Bottom: mass spectrum from the sample treated with the active enzyme, showing the formation of CDP-Rbo (m/z 536.06 ([M-H]-).

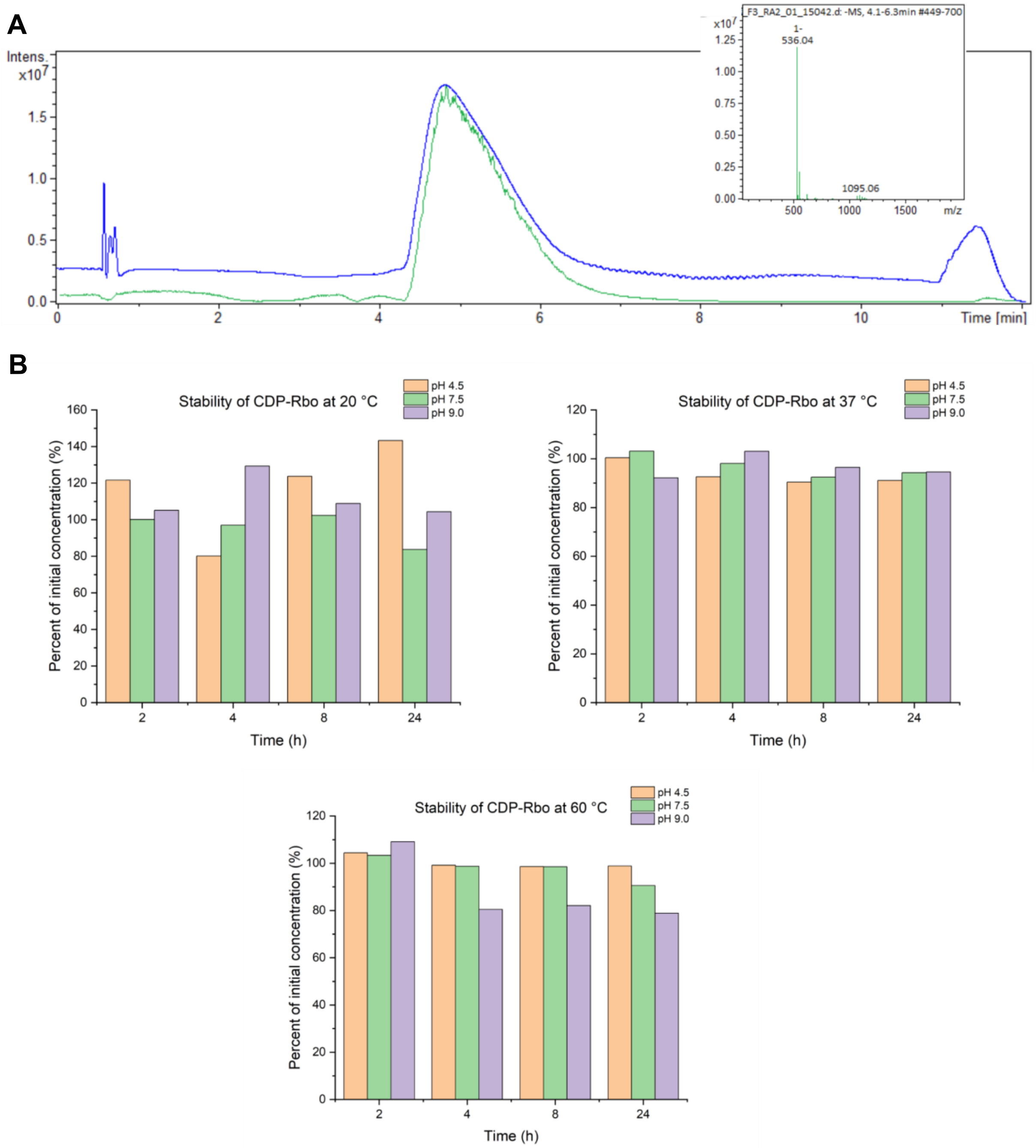


**Figure S3.** A) LC-MS chromatogram (green) and 280 nm UV trace (blue) of purified CDP-ribitol. The inset shows the mass spectrum corresponding to the peak, showing the expected mass of CDP-Rbo (m/z 536.04 ([M-H]⁻). B) Stability of CDP-Rbo during overnight incubation in buffers of different pH values and at different temperatures. Measurements were done using LC-MS/MS and quantified using a standard curve prepared from purified CDP-Rbo. Values were normalised to the concentration of CDP-Rbo at the start of the experiment.

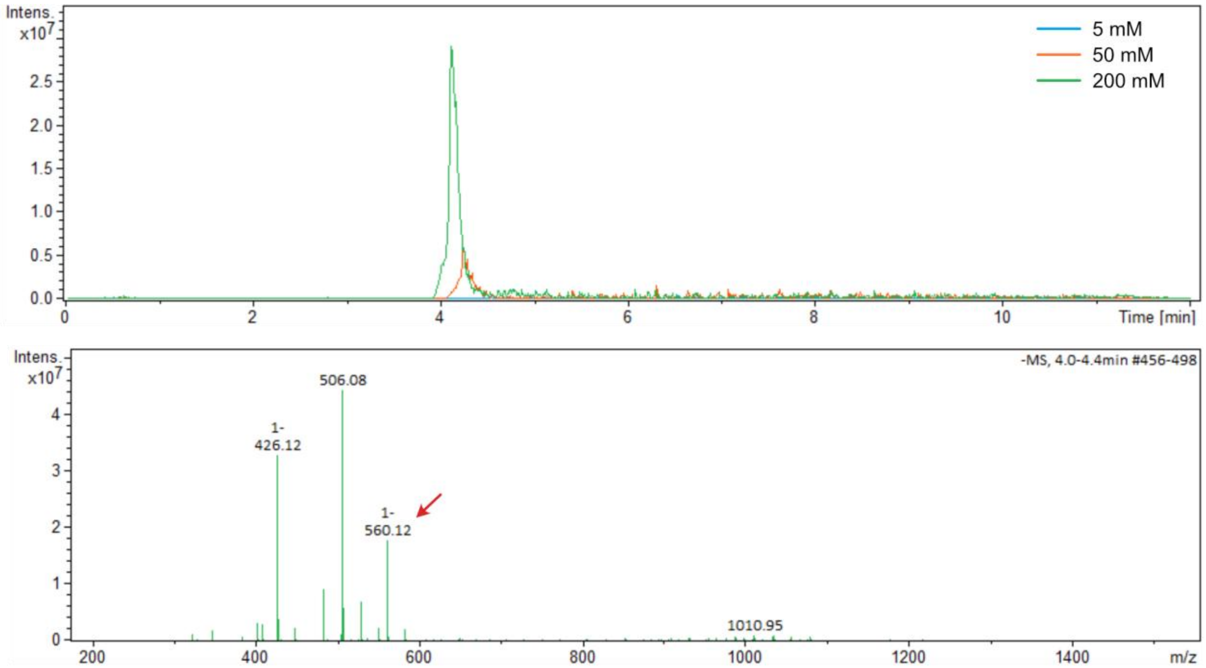


**Figure S4.** Top: Extracted ion chromatograms (EICs) of FGGY/TarI samples charged with different concentrations of probe A. The mass spectrum at the bottom corresponds to the sample with the highest concentration of the substrate, showing the conversion to its CDP-derivative (m/z 560.12 ([M-H]$^-$).

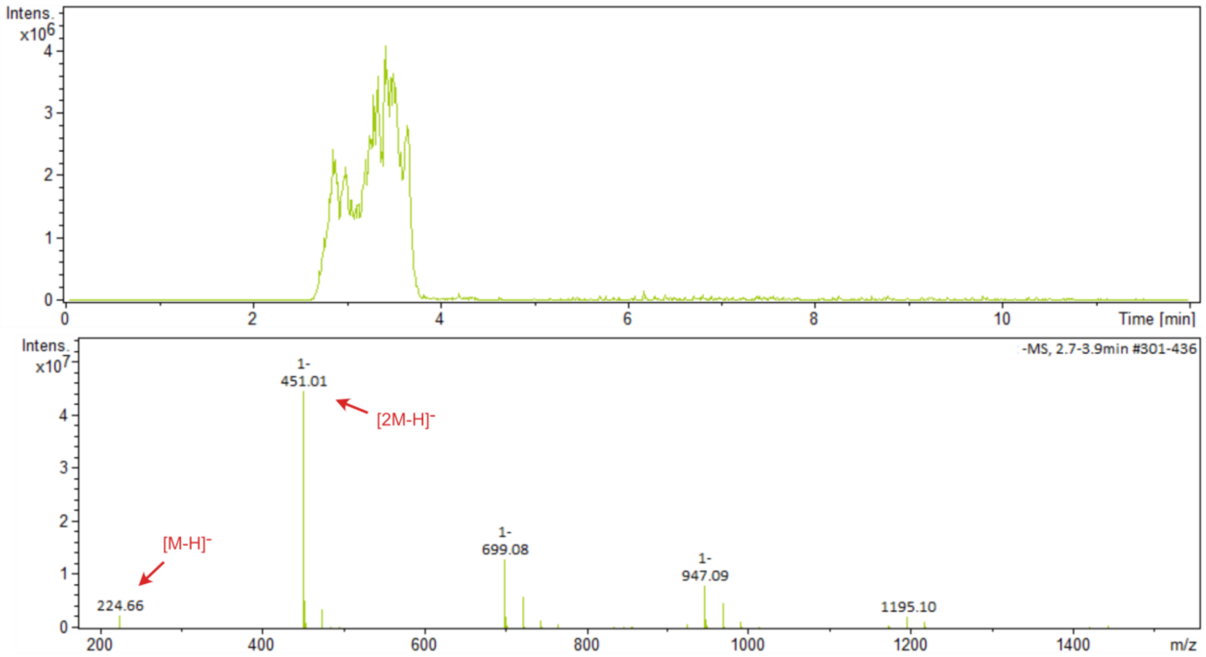


**Figure S5.** Top: Extracted ion chromatogram (EIC) of the phosphate derivative of probe B from a reaction with FGGY/TarI. Bottom: corresponding mass spectrum.

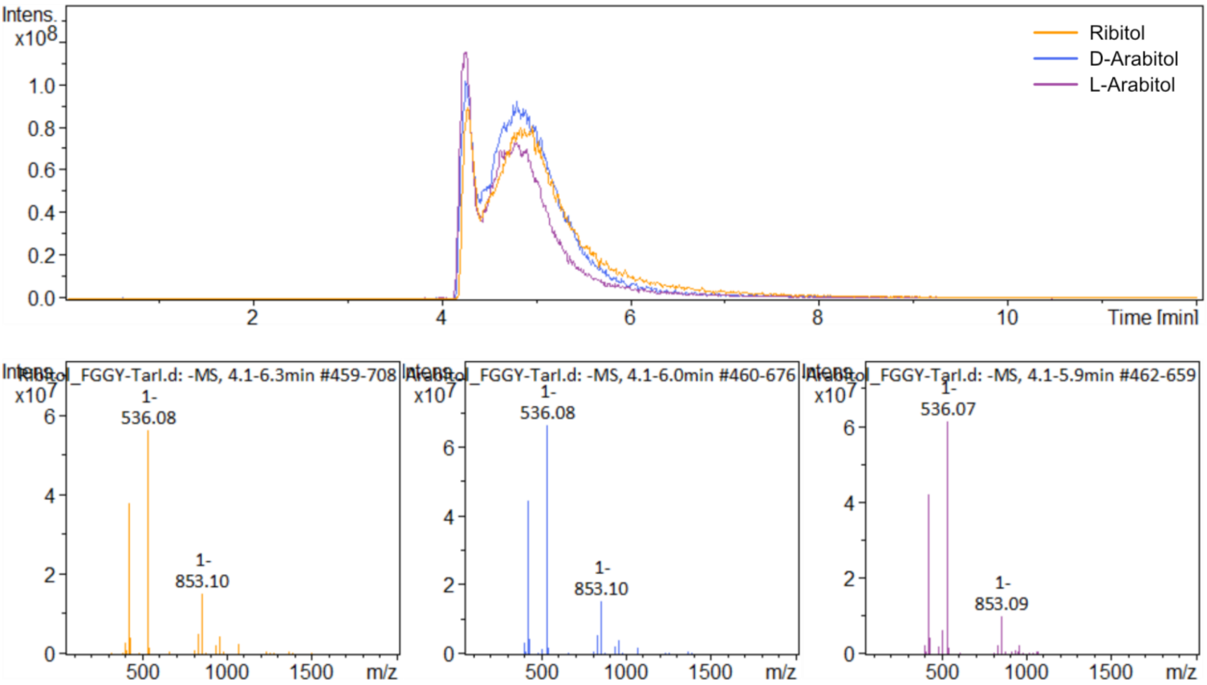


**Figure S6.** Top: Extracted ion chromatograms (EICs) of CDP-ribitol, CDP-D-arabinitol, and CDP-L-arabinitol from samples of their corresponding substrates treated with FGGY/TarI. Bottom: mass spectra of the CDP derivative for each substrate, showing the expected mass of the CDP conjugates (m/z 536.07/536.08 ($[M-H]^-$).

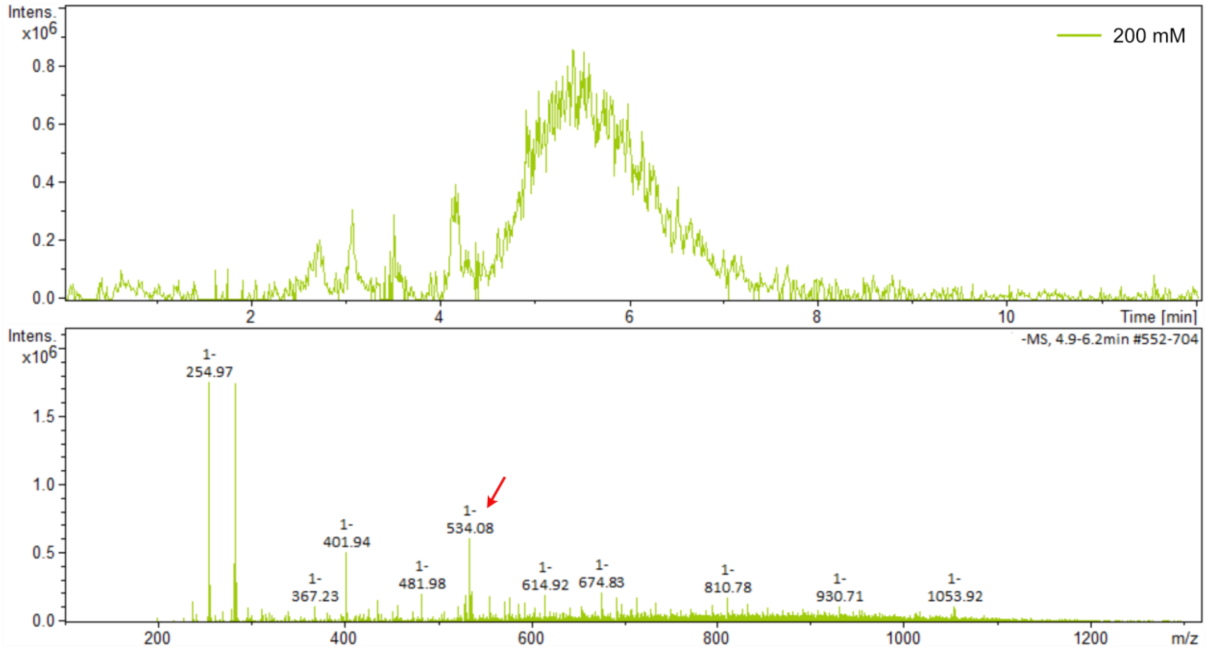


**Figure S7.** Top: Extracted ion chromatogram (EIC) of CDP-D-xylulose from a sample of the substrate charged with FGGY/TarI. The sample was treated with ALP to reduce the background noise from the free nucleotides. Bottom: corresponding mass spectrum, showing the expected mass of the product (m/z 534.08 ($[M-H]^-$).

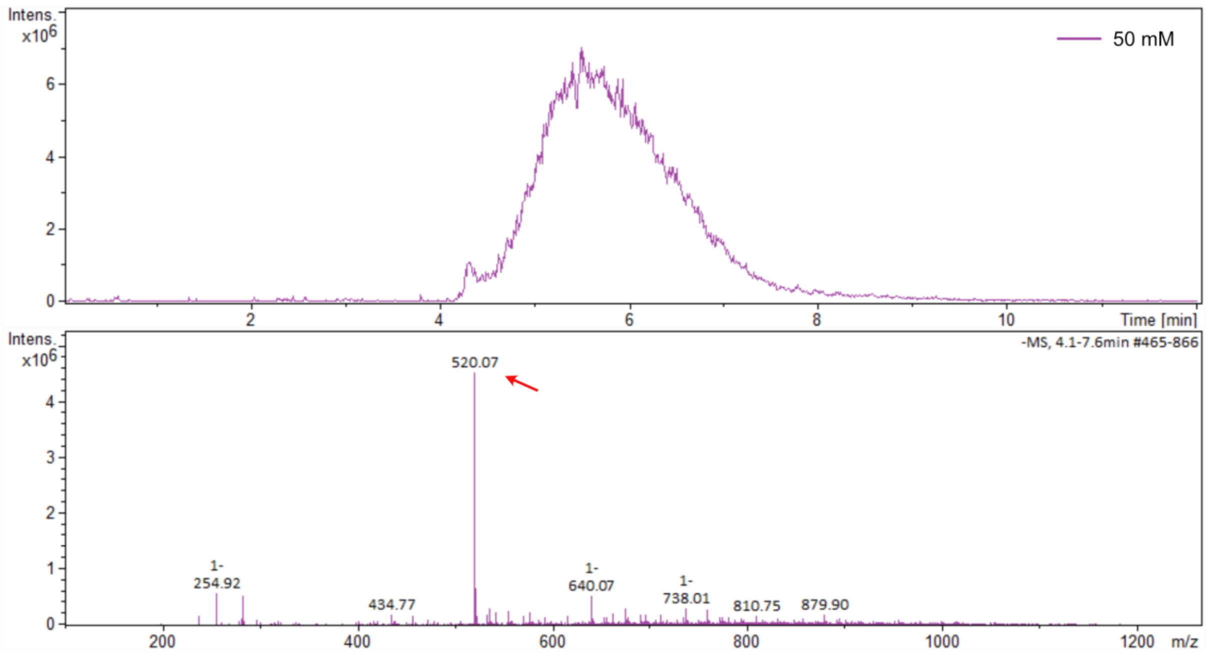


**Figure S8.** Top: Extracted ion chromatogram (EIC) of CDP-D-methyl erythritol from a sample of the substrate charged with FGGY/TarI. The sample was treated with ALP to reduce the background noise from the free nucleotides. Bottom: corresponding mass spectrum, showing the expected mass of the product (m/z 520.07 ([M-H]$^-$).

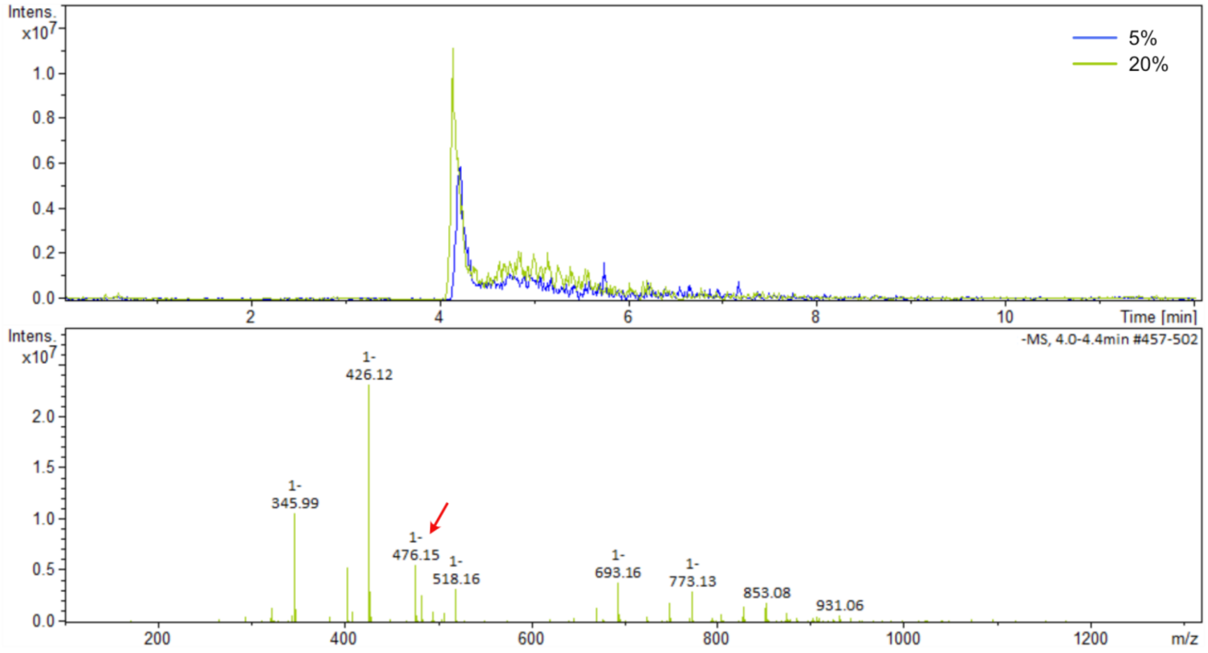


**Figure S9.** Top: Extracted ion chromatograms (EICs) of FGGY/PCYT2 samples charged with different concentrations of glycerol. The mass spectrum at the bottom corresponds to the sample with the highest concentration of the substrate, showing the conversion to its CDP-derivative (m/z 476.15 ([M-H]$^-$).

### NMR spectra

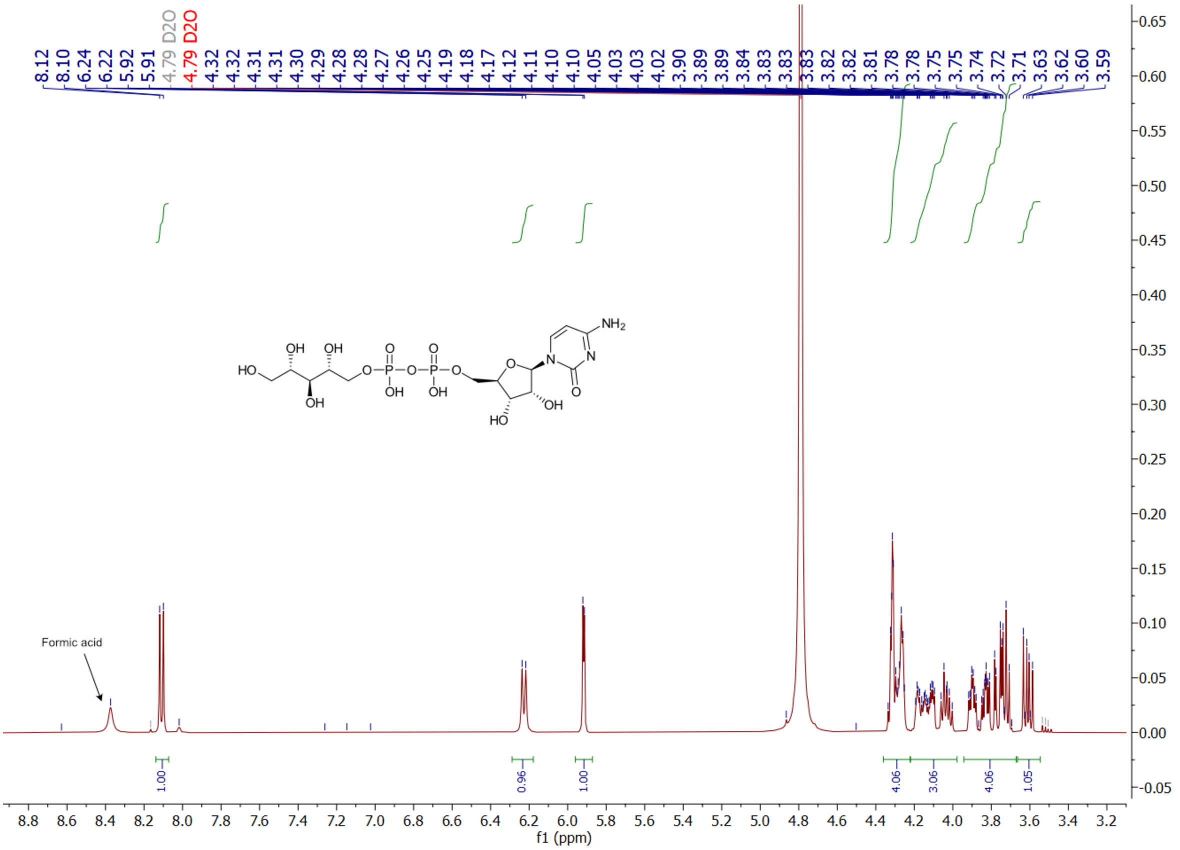
Formic acid
f1 (ppm)

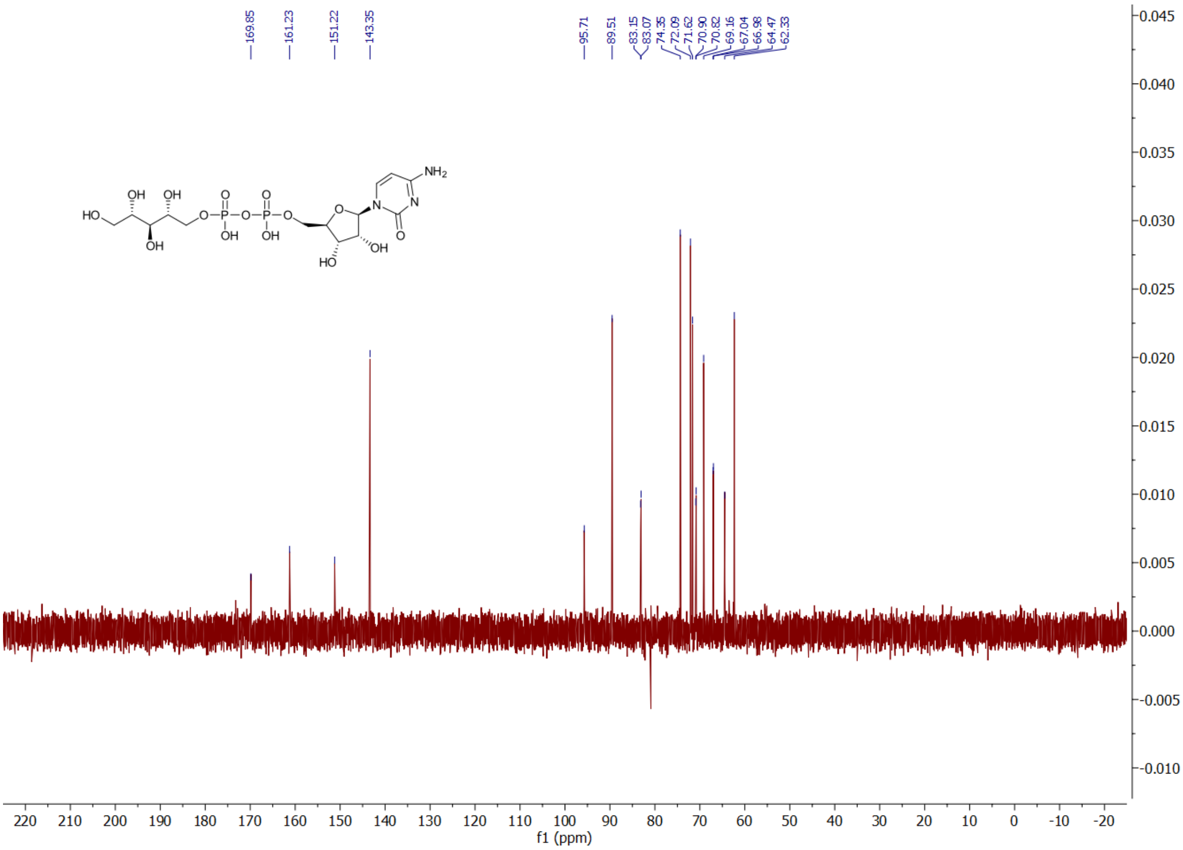
169.85
161.23
151.22
143.35
95.71
89.51
83.15
83.07
74.35
72.09
71.62
70.90
70.82
69.16
67.04
66.38
64.47
62.33
f1 (ppm)

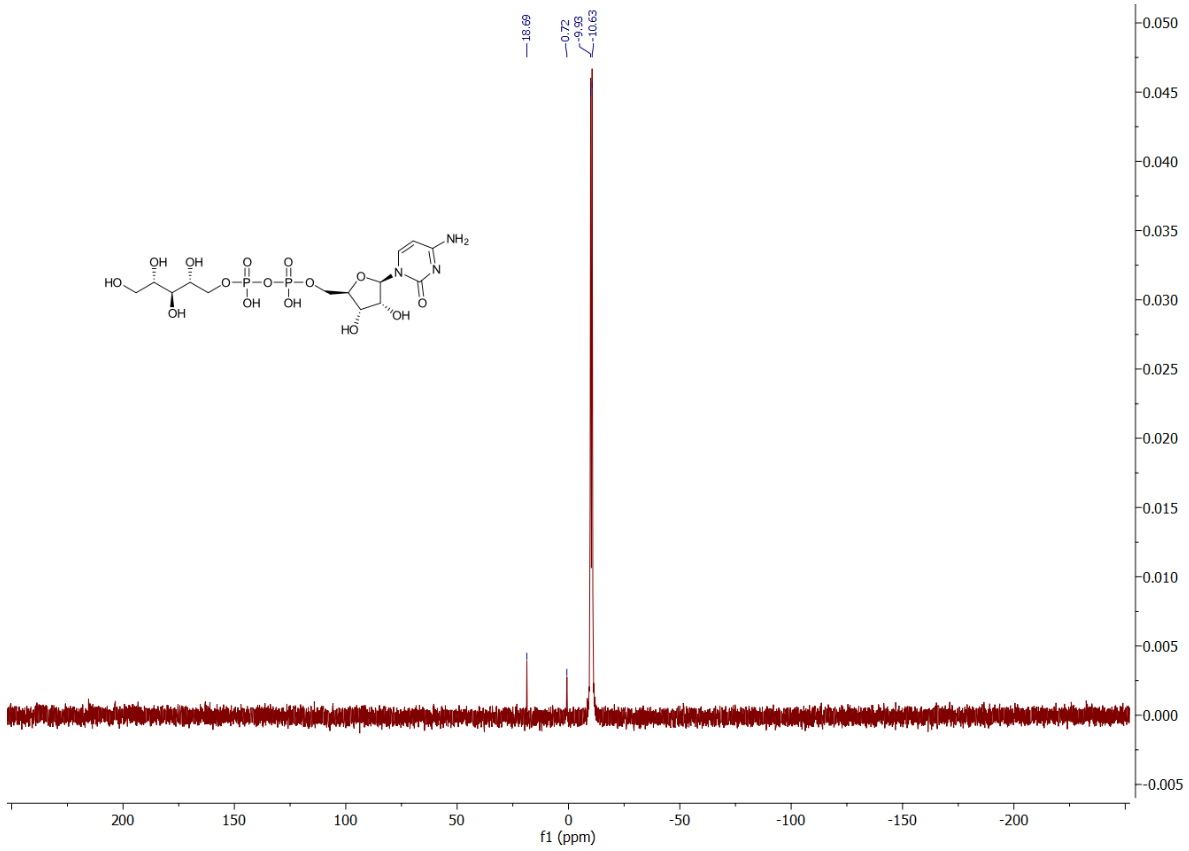
18.69
0.72
-9.93
-10.63
NH2
HO
OH
OH
OH
O
P
OH
OH
N
N
O
HO
f1 (ppm)
200
150
100
50
0
-50
-100
-150
-200
0.050
0.045
0.040
0.035
0.030
0.025
0.020
0.015
0.010
0.005
0.000
-0.005

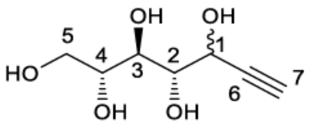
OH
OH
5
4
3
2
1
HO
6
7
OH
OH

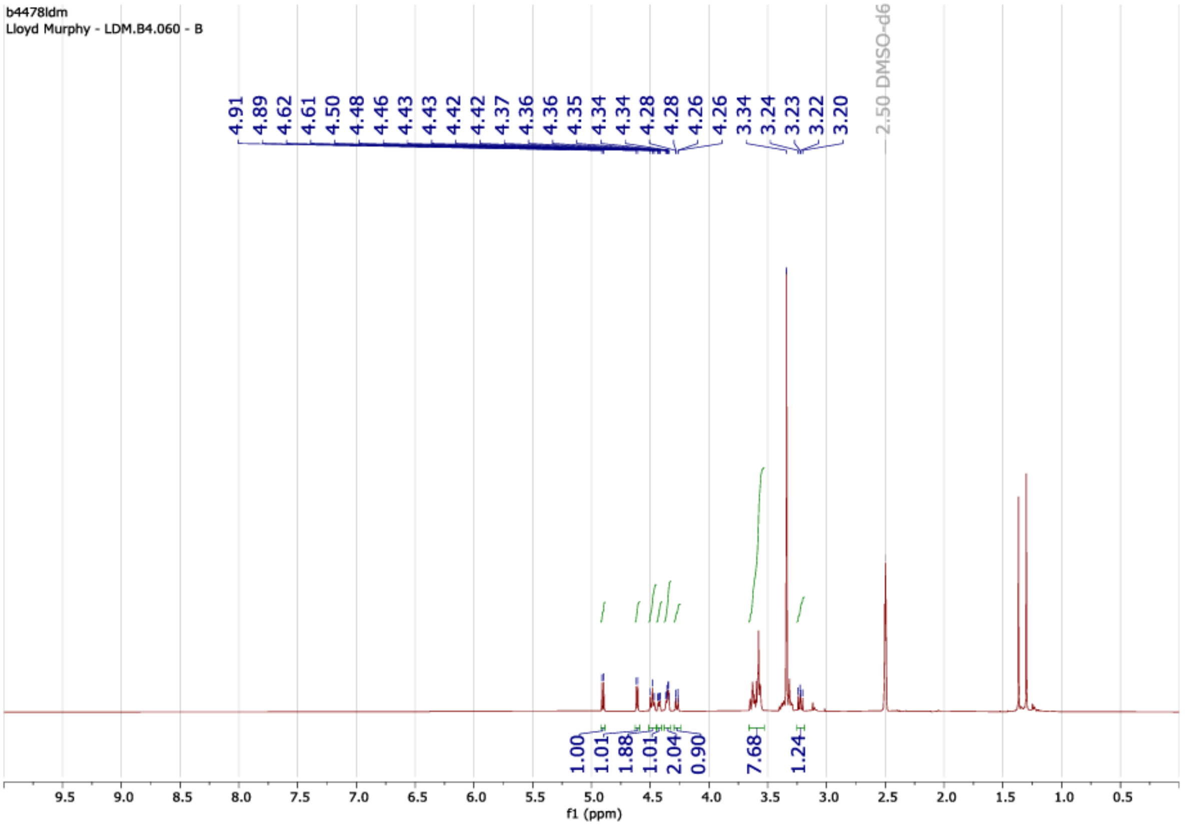
b4478ldm
Lloyd Murphy - LDM.B4.060 - B
4.91
4.89
4.62
4.61
4.50
4.48
4.46
4.43
4.43
4.42
4.42
4.37
4.36
4.36
4.35
4.34
4.34
4.28
4.28
4.26
4.26
3.34
3.24
3.23
3.22
3.20
2.50 DMSO-d6
1.00
1.01
1.88
1.01
2.04
0.90
7.68
1.24
f1 (ppm)

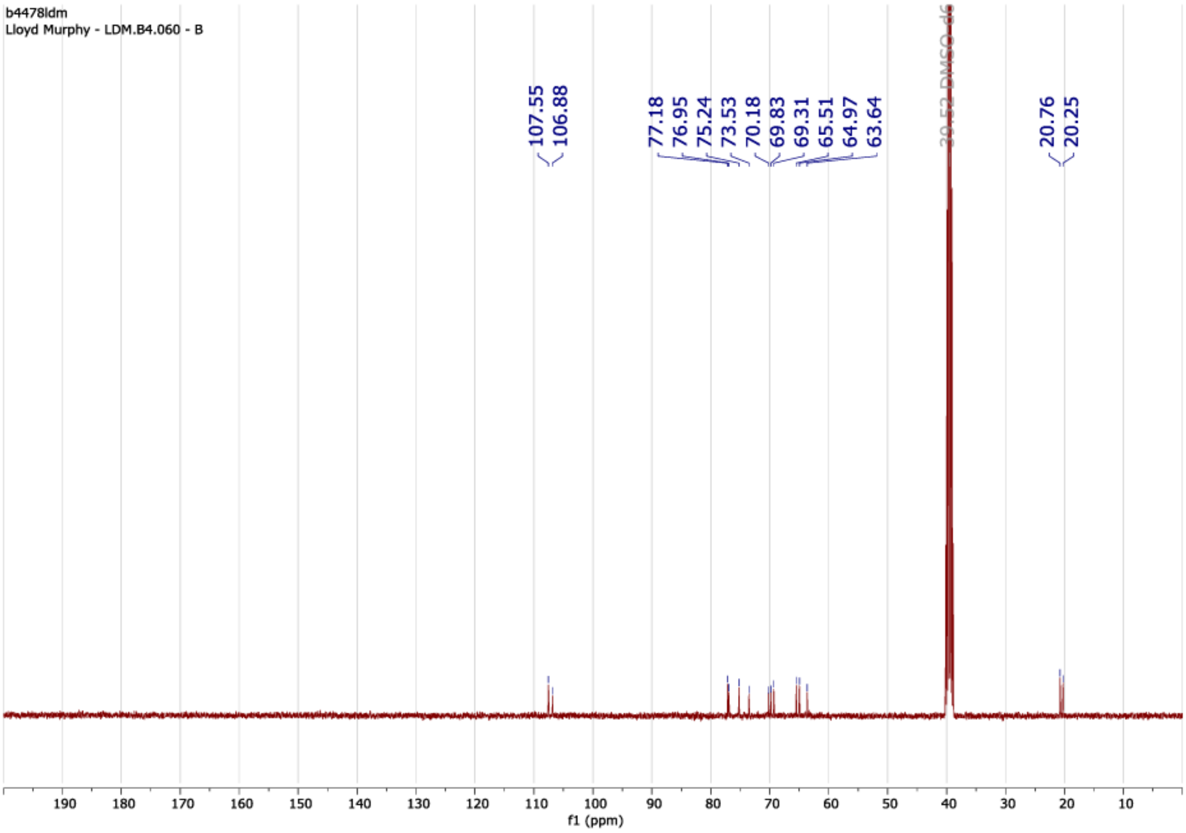
b4478ldm
Lloyd Murphy - LDM.B4.060 - B
107.55
106.88
77.18
76.95
75.24
73.53
70.18
69.83
69.31
65.51
64.97
63.64
39.52 DMSO-d6
20.76
20.25
f1 (ppm)

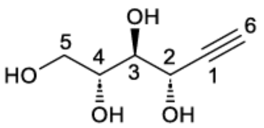
OH
6
5
HO
4
3
2
1
OH
OH

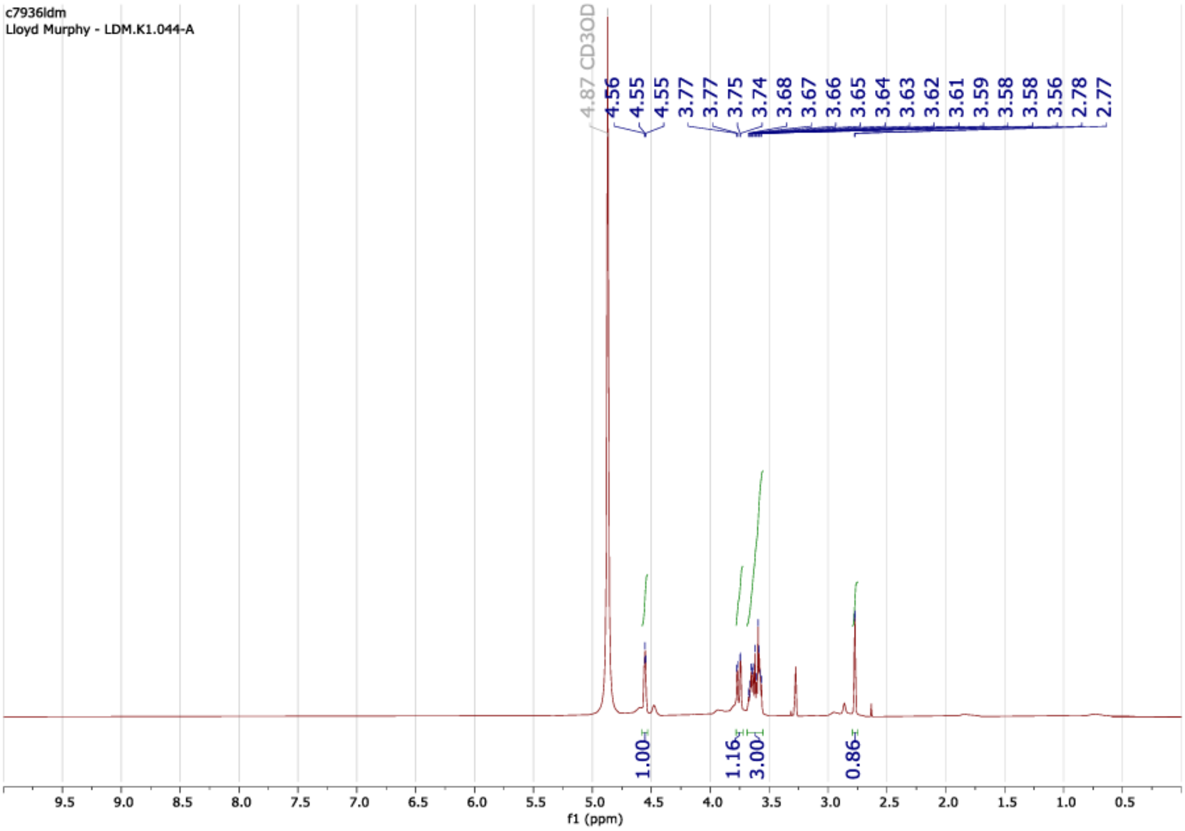
c7936ldm
Lloyd Murphy - LDM.K1.044-A
4.87 CD3OD
4.56
4.55
4.55
3.77
3.77
3.75
3.74
3.68
3.67
3.66
3.65
3.64
3.63
3.62
3.61
3.59
3.58
3.58
3.56
2.78
2.77
1.00
1.16
3.00
0.86
9.5
9.0
8.5
8.0
7.5
7.0
6.5
6.0
5.5
5.0
4.5
4.0
3.5
3.0
2.5
2.0
1.5
1.0
0.5
f1 (ppm)

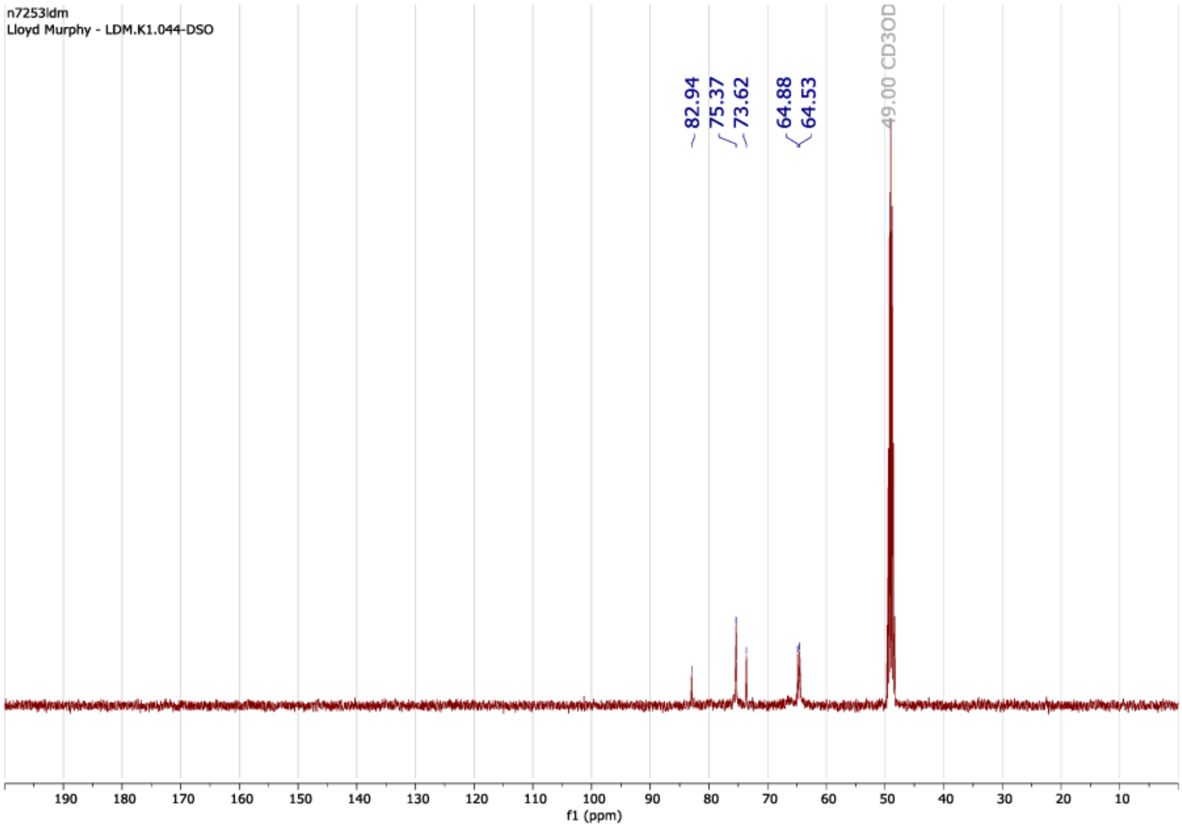
n7253ldm
Lloyd Murphy - LDM.K1.044-DSO
82.94
75.37
73.62
64.88
64.53
49.00 CD3OD
190
180
170
160
150
140
130
120
110
100
90
80
70
60
50
40
30
20
10
f1 (ppm)
